# Likely role of promoter reconstitution in Mpr-mediated D29 resistance by *Mycobacterium smegmatis*

**DOI:** 10.64898/2026.02.27.708680

**Authors:** Buhari Yusuf, Yanan Ju, Biao Zhou, Abdul Malik, Md Shah Alam, Lijie Li, Haftay Abraha Tadesse, Belachew Aweke Mulu, Cuiting Fang, Xirong Tian, Hongyi Chen, Li Wan, Liqiang Feng, Xiaoli Xiong, Shuai Wang, Tianyu Zhang

**Affiliations:** State Key Laboratory of Respiratory Disease, Guangzhou Institutes of Biomedicine and Health, Chinese Academy of Sciences, Guangzhou 510530, China; China-New Zealand Joint Laboratory on Biomedicine and Health, Guangzhou Institutes of Biomedicine and Health, Chinese Academy of Sciences, Guangzhou 510530, China; Guangdong-Hong Kong-Macao Joint Laboratory of Respiratory Infectious Diseases, Guangzhou Institutes of Biomedicine and Health, Chinese Academy of Sciences, Guangzhou 510530, China; University of Chinese Academy of Sciences, Beijing 100049, China; School of Life Sciences, University of Science and Technology of China, Hefei 230027, China; State Key Laboratory of Respiratory Disease, Guangzhou Chest Hospital, Institute of Tuberculosis, Guangzhou Medical University, Guangzhou 510095, China; Guangzhou Medical University-Guangzhou Institutes of Biomedicine and Health Joint School of Life Sciences, Guangzhou Medical University, Guangzhou 511436, China; State Key Laboratory of Respiratory Disease, Guangzhou Chest Hospital, Guangzhou, China; Guangzhou National Laboratory, Guangzhou 510005, China; West China School of Public Health, Sichuan University, Chengdu 610041, China

**Keywords:** *Mycobacterium smegmatis*, mycobacteriophage D29, promoter reconstitution, Mpr, phage resistance

## Abstract

The <u>m</u>ulti-copy <u>p</u>hage resistance gene (*mpr*) of *M. smegmatis* is a major factor in resistance to the lytic mycobacteriophage D29. Mpr is a membrane-bound exonuclease that cleaves phage DNA post injection, hence blocking downstream stages in the phage infection cycle. The mechanism of resistance allows for adsorption, is non-abortive and independent of any mutation in the gene. Rather, it depends on overexpression of a wild type copy of the gene. However, the underlying factor behind *mpr* overexpression in spontaneous D29-resistant mutants of *M. smegmatis* remained elusive. Here, we report that D29 infection triggers insertion sequence (IS) rearrangements, including the transposition and integration of IS6120 directly upstream of *mpr*. Mutants with IS6120 integration upstream of *mpr* show highly elevated Mpr expression. Whole genome sequence analysis reveals that IS6120 introduced a putative transcription factor-binding site and a canonical -35 promoter element at the integration site, hence reconstituting a fuller promoter (*rcp*) than the original promoter (*wtp*) at the site. Promoter reporter assays suggest that *rcp* is a far stronger promoter than *wtp*, implying that elevated *mpr* expression in D29-resistant mutants with this transposition event could be due to promoter reconstitution. While strains with this transposition event appear to grow normally, *rcp*-driven, vector-borne Mpr overexpression appears to be toxic as it barely allows for colony formation on agar plates. This study reports a previously unknown factor likely behind *mpr* regulation in *M. smegmatis*, adding to the existing knowledge of mycobacterial anti-phage defense mechanisms and guiding rational phage engineering efforts for therapeutic applications.

## Introduction

Antimicrobial resistance (AMR) remains a serious public health threat, and despite efforts to develop novel therapeutics or vaccines, the threat remains ever existent. This is because the pace at which bacterial pathogens develop resistance to antimicrobials surpasses the rate at which new therapeutics are introduced into clinical use. Conventional antibiotic treatments occasionally fail due to extensive drug resistance, hence complicating disease management. In fact, it is estimated that global death toll due to bacterial AMR may surpass 39 million between 2025 and 2050 (1). For this reason, interest in phage therapy as an alternative to conventional antibiotic therapy is gaining significant traction.

Phage therapy is the use of bacteria-eating viruses (or bacteriophages) to treat bacterial infections. Bacteriophages are highly specific, relatively safe and hold significant potential as alternatives to redundant, toxic antibiotic therapy. Currently, phage therapy is only available to patients under the framework of compassionate use (2). Compassionate use of phage therapy against drug-resistant mycobacterial infections have further highlighted the significant potential of phage therapy even in complicated cases of failed antibiotic therapy (3–9). However, bacterial pathogens can develop resistance to bacteriophages, which represents a potent threat to sustainable utility of phage therapy.

Bacterial pathogens have evolved an extensive arsenal of anti-phage defense systems, including clinical bacterial isolates (10–13). The lytic mycobacteriophage D29 has demonstrated therapeutic potential in compassionate use against BCG and *Mycobacterium abscessus* infections (5,7). However, a resistance mechanism against D29 and its parent L5 already exists in the model bacterium *Mycobacterium smegmatis* (14). The <u>m</u>ulticopy <u>p</u>hage resistance gene (*mpr*) of *M. smegmatis* is a major factor in D29 resistance. Mpr is a membrane-bound exonuclease that mediates resistance via cleavage of phage DNA post injection, which blocks downstream stages of the phage lifecycle and inhibit phage propagation (15). Mpr contains a domain of unknown function (DUF4352) that is believed to be of viral origin, suggesting that *M. smegmatis* obtained this protein during the course of its evolution with viral predators and repurposed it for anti-phage defense (16). The protein has also been reported to aid non-homologous end joining (NHEJ)-mediated anti-phage defense in *M. smegmatis* (17), further highlighting its relevance to antiviral defense. However, the role of Mpr in D29 resistance is independent of any genetic change in the gene, but rather depends on overexpression of a wild type copy of the gene. The exact factor behind elevated Mpr expression in spontaneous D29-resistant mutants remained unknown.

We report here that infection with D29 triggers IS rearrangements in *M. smegmatis*, including the transposition and integration of IS6120 directly upstream of *mpr*. IS6120 introduces a canonical - 35 promoter element as well as a putative transcription factor-binding site (TFBS) for leucine-responsive regulatory protein (Lrp), hence reconstituting a fuller promoter than the original promoter at the integration site. Promoter reporter assays show that the reconstituted promoter (*rcp*) is far stronger than the wild type promoter (*wtp*), which could represent a previously unknown regulatory mechanism for the toxic protein Mpr in *M. smegmatis*.

## Materials and Methods

### Bacterial strains, phage and growth conditions

Cloning experiments were carried out in *E. coli* Trans1 (TransGen Biotech., China). *E. coli* was maintained in liquid or on solid Luria-Bertani medium. Selective growth of *E. coli* was carried out in the presence of kanamycin (kan, 50 µg/mL), ampicillin (amp, 100 µg/mL) or zeocin (zeo, 30 µg/mL). *M. smegmatis* mc^2^ 155 was maintained in Middlebrook 7H9 (Difco) medium supplemented 10% OADC, 0.2 % glycerol and 0.05% Tween 80. *M. smegmatis* was selectively grown in liquid or on solid medium in the presence of the following concentrations of antibiotics: kan (liquid, 20 µg/mL; solid, 50 µg/mL); zeo (30 µg/mL). All liquid cultures were maintained at 37 °C with shaking at 200 revolutions per minute (rpm). The phage used in this study is the lytic mycobacteriophage D29. All D29 infection experiments were carried out at 37 °C in the presence of 2 mM CaCl_2_.

### Phage amplification

Wild type *M. smegmatis* (Ms^Wt^) was grown in 7H9 medium to OD_600_ 0.8-1 at 37 °C with shaking. The cells were pelleted and washed thrice in MP buffer, then re-suspended in 1.5 mL of the same buffer. Ten-fold serial dilutions of the phage stock were prepared in MP buffer until 10^-4^ dilution factor. A four-component mixture was prepared by adding top agar, washed Ms^Wt^ cells, D29 at 10^-4^ dilution and CaCl_2_ at a final concentration of 2 mM. The mixture was mixed by pipetting and poured into plain 7H10 plates, spread evenly, allowed to dry and incubated at 37 °C for 12-24 hours. This was followed by addition of 2.5 mL of sterile MP buffer into each plate and rocking on an orbital shaker at room temperature for 4-5 hours. The MP buffer was then recovered from the plates by pipetting, centrifuged at 3,500 ×g for 10 minutes, then passing the supernatant through a 0.22 µm filter. The filtrate containing the amplified phage was stored at 4°C.

### Generation of spontaneous D29-resistant strains

Spontaneous D29-resistant strains were generated by subjecting Ms^Wt^ to four sequential rounds of extended exposures to mycobacteriophage D29 to increase the chance of retaining resistant clones and eliminating susceptible ones (18,19). In the first exposure cycle, bacterial cells were grown to OD_600_ 1.0, washed three times in MP buffer and re-suspended in 1 mL of MP buffer, infected with D29 and incubated at 37 °C for 7 hours with regular shaking. Cells from the 7-hour incubation were collected by centrifugation at 13,000 ×g for 3 minutes, washed three times in PBS-Tween 80, and re-suspended in 1 mL of the same medium. 50 uL of the washed cells were sub-cultured in 7H9 medium containing 0.05 % Tween 80 until OD_600_ 1.0. This resulting culture was washed three times in MP buffer, infected with D29 and incubated at 37 °C for 14 hours with regular shaking. This continued until the fourth exposure cycle, with the incubation period extended by 7 hours after the previous exposure cycle (21 and 28 hours for third and fourth exposure cycles, respectively). For every infection cycle, MP buffer was used as the medium of infection, bacterial cells were infected at MOI of 5, and it was ensured that CaCl_2_ was added to a final concentration of 2 mM. After the final 28-hour exposure, the cells were collected by centrifugation, washed in PBS-Tween 80, diluted and plated to collect single colonies.

### Phage susceptibility testing (PST)

Phage susceptibility testing was carried out by plaque and spot-kill assays as well as on D29-seeded 7H10 plates. Bacterial cells were grown in 7H9 medium to OD_600_ 0.8-1. The grown cells were pelleted by centrifugation at 3,500 ×g for 10 minutes. The supernatant was discarded and the pellet was washed three times in MP buffer and re-suspended in 0.5-1 mL of the same buffer. Ten-fold serial dilutions of D29 were prepared. For plaque assay, a soft overlay of top agar, phage, bacterial cells and 2 mM CaCl_2_ was prepared and poured into plain 7H10 plates in triplicates. The plates were dried and incubated at 37 °C for 12-24 hours before enumeration of plaque forming units. For spot-kill assay, a soft overlay of top agar, bacterial cells and CaCl_2_ was poured into a 7H10 plate, spread evenly and allowed to dry. Two microliters of ten-fold serial dilutions of the phage were spotted on the dried soft overlay, allowed to dry, and incubated at 37 °C for 12-24 hours. For PST on phage-seeded plates, 2 µL of ten-fold serial dilutions of the bacteria was spotted on 7H10 plate seeded with 10^7^ – 10^9^ plaque forming units per milliliter (PFU/mL) of D29, allowed to dry, and incubated at 37 °C for 3-5 days. Serial dilution of the bacteria was carried out in MP buffer containing 2 mM CaCl_2_, and 7H10 plate containing no phage was used as a negative control.

### Phage adsorption assay

Phage adsorption assay was carried out as described previously (19,20). Bacterial cells were grown to mid log phase, washed three times and re-suspended in MP buffer. The cells were diluted in MP buffer to OD_600_ 0.3. Ten-fold serial dilutions of D29 were prepared in MP buffer until 10^-3^. To determine terminal phage adsorption, 30 µL of 10^-3^ dilution of the phage was added to 200 µL aliquot of diluted bacteria, and CaCl_2_ was added to a final concentration of 2 mM. The mixture was incubated at 37 °C with shaking for 60 minutes. Simultaneously, same quantity of same phage inoculum was added to 200 µL of sterile MP buffer, 2 mM CaCl_2_ was added, mixed, and immediately centrifuged at 5000 ×g for 3 minutes. The supernatant was collected, diluted and used for plaque assay on MP buffer-washed Ms^Wt^. After the 60 minutes shaking of phage-bacteria mixture, the mixture was also centrifuged at 5000 ×g for 3 minutes, the supernatant was collected, diluted and used for plaque assay on MP buffer-washed Ms^Wt^. Plates were allowed to dry, incubated at 37 °C for 24 hours before enumeration of plaque forming units. Plates were prepared in triplicates, and data was presented as percentage of adsorbed phage. For determination of phage adsorption rates, same phage-bacteria mixtures and incubation conditions were prepared for plaque assay at 0-, 30- and 60-minute time points. Collected data was presented as percentage of unadsorbed phage.

### Colony morphology and spot inhibition by kanamycin and zeocin

All strains were grown in 7H9 medium to OD_600_ 0.8-1 and diluted to OD_600_ 0.6. Two microliters from diluted culture of each strain was spotted on plain 7H10 plate and allowed to dry. For susceptibility testing to kan (50 µg/mL) or zeo (30 µg/mL), the cells were spotted on 7H10 plates containing indicated concentration of either drug. Dried plates were incubated at 37 °C for 3-4 days.

### Whole genome sequencing (WGS)

Genomic DNA (gDNA) extraction and sequencing were carried out by Shanghai Jingnuo Biotechnology Co., Ltd. (Shanghai, China). A modified Cetyltrimethylammonium bromide (CTAB) method was used for extraction of gDNA (21). Extracted gDNA was quantified using Qubit 2.0 Fluorometer (ThermoFisher, USA), followed by library construction using Hieff NGS^®^ OnePot Flash DNA Library Prep Kit (Yeasen, China) by enzymatic fragmentation of gDNA, end repair, tail adenylation, adapter ligation, purification and PCR amplification. The library was initially quantified using Qubit 2.0 and diluted. Fragment sizes were verified using Agilent 2100 bioanalyzer (Agilent, USA) and effective library concentration was quantified again by qPCR. Libraries were pooled onto a flow cell, clustered using cBOT and sequenced using the NovaSeq 6000 high-throughput sequencing platform (Illumina, Inc., USA).

### Bioinformatics

Next generation sequencing (NGS) data was analyzed on the GalaxyEU bioinformatics server using default settings (22). Raw sequence reads were checked for quality using FASTQC (23), and undesirable reads/reads of low quality were trimmed using FASTP (24). Mapping to the reference genome (*M. smegmatis* mc^2^ 155, NCBI accession number CP000480.1) and variant calling were achieved using Snippy (25), and the output data was exported as VCF and tab-separated files. All detected variants were verified by Sanger sequencing.

### RNA extraction and RT-qPCR

RNA extraction was carried out according to manufacturer instructions using Magen HiPure Bacterial RNA Mini Kit (Magen, China). For bacterial growth to detect mRNA levels in the presence and absence of D29 infection, the strains were first washed thrice in 7H9 medium without Tween 80, sub-cultured in 7H9 (without Tween 80) containing 2 mM CaCl_2_ with/without D29 at MOI 5. The strains were then washed thrice in PBS-Tween 80 prior to RNA extraction. Extracted RNA was reverse transcribed using ExonScript All-in-One RT SuperMix with DNase (EXONGEN, China) according to manufacturer instructions. The RT-qPCR reaction was prepared and run according to manufacturer instructions using Taq Pro Universal SYBR qPCR Master Mix (Vazyme Biotech., China). All reactions were run in C1000 Touch^TM^ Thermal Cycler (Bio-Rad, USA). SigA was used as control, and all wild type/untreated group expressions were normalized to 1, and the treated/test group expression was presented as fold change. Expression levels (fold) were represented as bar graphs using GraphPad Prism version 9.0.0 (GraphPad, USA).

### IS transposition analysis

Sequences of the different IS elements encoded by *M. smegmatis* genome were obtained from ISFinder database (26). ISMapper (27) was used to map out the different IS integration sites in raw sequence reads of each sequenced D29-resistant mutant. Briefly, paired end Illumina reads of a D29-resistant mutant, a FASTA file of the IS query sequence and a genbank file of the reference genome (*M. smegmatis* mc^2^ 155, NCBI accession number CP000480.1) were used as input files. The paired end Illumina reads were mapped to the IS query sequence using BWA-MEM (28), which generates a SAM file as output. Sequences directly flanking the IS query (i.e. pairs of unmapped reads that mapped to the end of the IS query sequence) as well as soft clipped regions (reads containing sequences that mapped to the IS query and extend into neighboring sequence) were extracted from the SAM file using SAMtools (29) and SAMBLASTER (30) respectively. The extracted sequences were converted to FASTQ format using BEDTools (31) and filtered using BioPython to extract soft clipped read portions that are 5-30 bp in length. The resulting sequences were sorted into left and right flanking sequences and separately mapped to the reference genome using BWA-MEM, and BEDTools was used to determine the exact position of IS integration in the D29-resistant mutant. All detected IS integrations were verified by PCR.

### Promoter reporter assay

#### Lawn appearance

Strains were grown in 7H9 medium to OD_600_ 0.8-1, then diluted to OD_600_ 0.6. Ten microliters from diluted culture of each strain was streaked on a plain 7H10 plate. The plate was dried and incubated at 37 °C until appearance of clear bacterial lawn.

#### Live fluorescence

Strains were grown in 7H9 medium to OD_600_ 0.8-1, then diluted to OD_600_ 0.6. Two-fold serial dilutions of the diluted cultures were prepared in triplicates for each strain in a 96-well microplate to obtain OD_600_ 0.6, 0.3, 0.15, 0.075, 0.0375 and 0.01875. Fluorescence was quantified in FlexStation 3 Multi-Mode Microplate Reader (Molecular Devices, USA) at excitation and emission wavelengths of 488 nm and 520 nm respectively. Quantification data for fluorescent strains were normalized by subtracting Ms^Wt^ fluorescence from that of each strain, and the results were plotted as bar graphs in GraphPad Prism.

#### Flow cytometry

Flow cytometry was carried out as previously described (32) with modification. The strains were grown in 7H9 medium to OD_600_ 0.8-1. Two milliliters of each strain was pelleted, washed twice in PBS-Tween 80, and fixed in 1 mL of 4 % glutaraldehyde in PBS-Tween 80 at room temperature for 30 minutes. The fixed cells were pelleted, re-suspended in 2 mL of sterile PBS, and passed through a 40 µm filter prior to flow cytometry using a CytoFLEX flow cytometer (Beckman Coulter, USA). A minimum of 10,000 cells were selected to count in focus GFP events for each strain. Flow cytometry data was analyzed using FlowJo version 10.8.1.

#### Confocal microscopy

Strains were grown in 7H9 medium to OD_600_ 0.8-1. One milliliter of each strain was pelleted and washed twice in PBS-Tween 80. The supernatant was discarded, the cells were re-suspended in 1 mL of 4 % glutaraldehyde in PBS-Tween 80 and allowed to stand at room temperature for 30 minutes. The fixed cells were pelleted and re-suspended in 200 µL of PBS-Tween 80. For each strain, 5 µL of re-suspended fixed cells as well as a 10-fold serial dilution were mounted on a microscope slide and covered with a coverslip, then observed under a Zeiss LSM 800 Confocal Laser Scanning Microscope (Zeiss, Germany).

#### Drug susceptibility testing (DST)

Drug susceptibility testing was carried out by broth microdilution. Bacterial strains were cultured in 7H9 medium at 37 °C with shaking until OD_600_ 0.8-1, then diluted to a cell density of 10^5^ colony forming units per milliliter (CFU/mL) in 7H9 without Tween 80. Bacterial cultures in two-fold serial dilutions of each drug as well as a plain control were prepared in 96-well microplates. The plates were incubated at 37 °C for 3 – 5 days. Minimum inhibitory concentration (MIC) was defined as the concentration at which visible bacterial growth was inhibited.

## Results

### Spontaneous D29 resistance in *M. smegmatis*

We screened 137 colonies of *M. smegmatis* for D29 resistance after multiple cycles of exposure, and a vast majority of them (82.5 %, n = 113) were classified as D29-resistant isolates. We then selected and purified 24 colonies, re-screened them for D29 resistance and sequenced their genomes. Of these 24 isolates, two strains, A72.1 and 2.6.1, particularly stand out for their high level of resistance to D29. Both strains appear to manifest no obvious change in surface morphology, but exhibit high level of resistance when screened by spot-kill and plaque assays (Figure 1A, B), which suggests impaired D29 infection efficiency against A72.1 and 2.6.1. Notably, greater susceptibility is seen at higher D29 concentration (Figure 1A), possibly suggesting dose-dependent resistance. Phage adsorption assay indicates little to no effect on D29 adsorption to both strains, suggesting that the mechanism of resistance in both strains occurs at stage(s) beyond phage adsorption (Figure 1C). The D29-resistant mutants A72.1 and 2.6.1 sustained point mutations and two IS transposition events each, including IS6120 integration directly upstream of *mpr* in both mutants.

**Figure 1.**
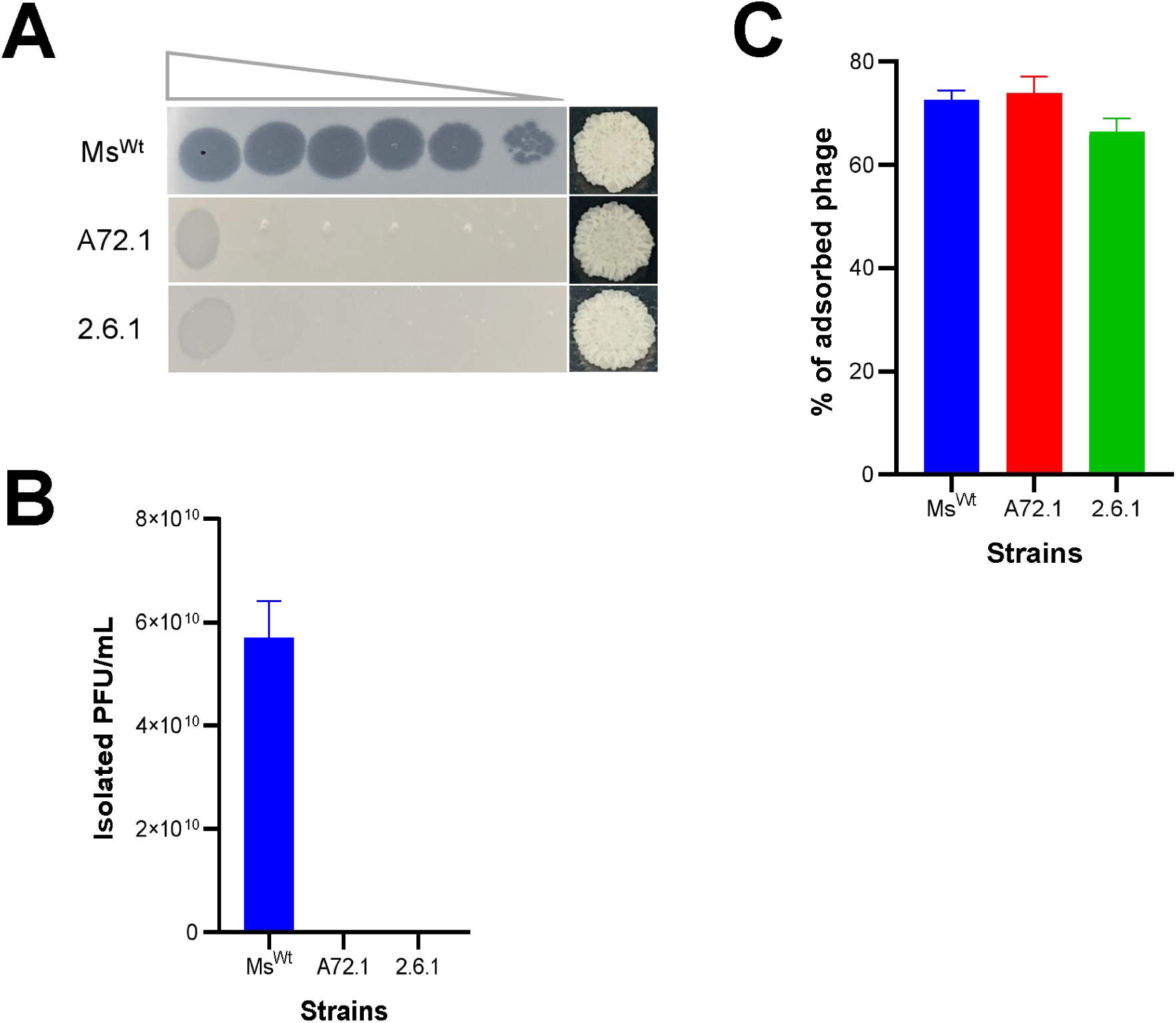
Spontaneous D29 resistance in *M. smegmatis.* A, A72.1 and 2.6.1 were tested for D29 resistance by spot-kill assay, and both appear to be particularly highly resistant to D29 compared to Ms^Wt^. B, both strains were also tested for D29 resistance by plaque assay, which appears to be consistent with the spot-kill assay. C, terminal adsorption suggests that D29 adsorption is not inhibited in the two strains, suggesting that the resistance mechanism is at a stage downstream of phage adsorption.

### D29 infection triggers IS rearrangements in *M. smegmatis*

It is likely that other genetic factors alone or in combination with the identified mutations may be responsible for D29 resistance phenotypes of the mutants (20). For this reason, we carried out IS transposition analysis on raw FASTQ data obtained for each of the mutants as well as Ms^Wt^. We first collected the sequences of all known IS elements encoded by the *M. smegmatis* genome from ISFinder database (26) and determined their prevalence across the genome (Table S1, Figure S1). We then identified all IS rearrangements that were present in the mutants but not in Ms^Wt^, then verified them by PCR alongside the Ms^Wt^ control (Figure 2).

**Figure 2.**
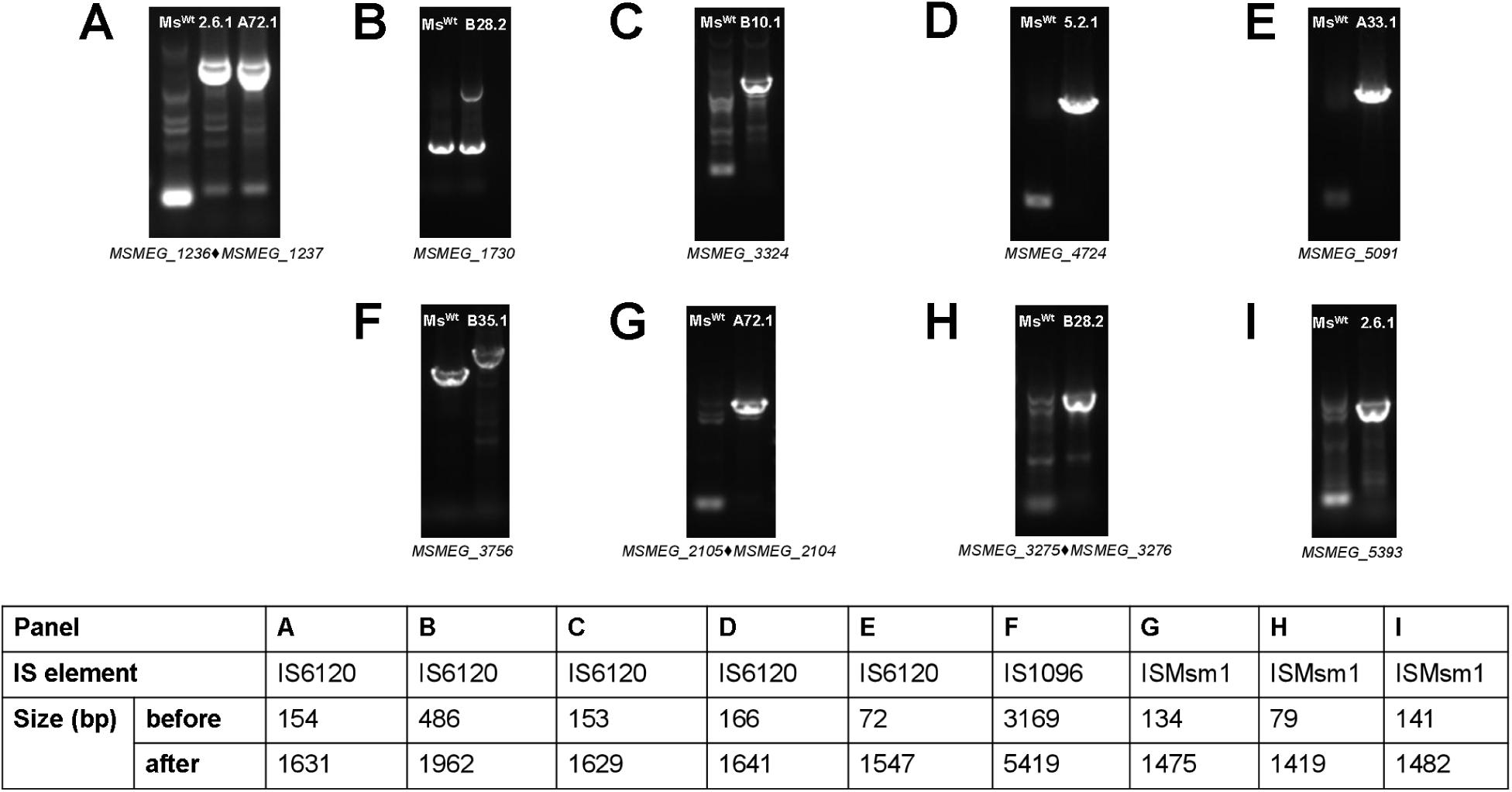
PCR verification of IS rearrangements. A total of 7 (29.2 %) of the sequenced D29-resistant mutants sustained at least a single IS transposition event. PCR verification of integration at each site as well as affected gene/region are shown in the upper panels A-I, whereas sizes at the site before and after IS integration are outlined in the lower table in the figure. Integration at intergenic region is indicated by a “black diamond icon” between the two genes.

A total of 10 IS transposition events were detected in seven D29-resistant mutants, where 4 mutants had single event each and 3 had double. There had been six IS6120, three ISMsm1 and one IS1096 transposition events in the mutants, in which the IS integrations took place in six coding and three non-coding regions. A72.1 and 2.6.1 – the two strains that manifested incredibly higher level of D29 resistance compared to the other mutants – appear to have sustained two IS transposition events each, making them strains of particular interest (Figure 3). Details about the integration position and protein products of the affected genes are outlined in Table S2.

**Figure 3.**
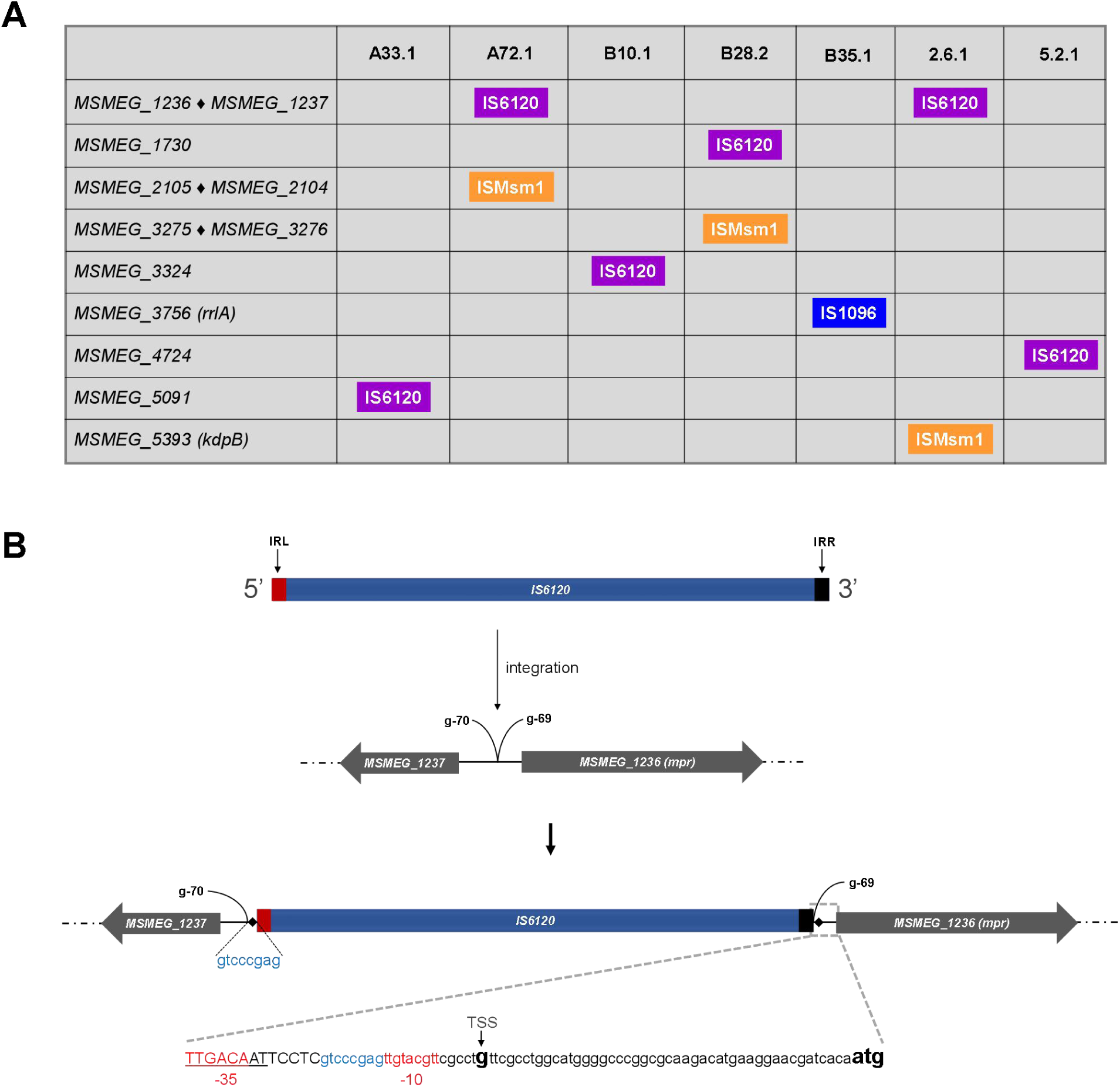
IS rearrangements in the D29-resistant mutants. A, the different strains/regions affected by IS rearrangements. IS integration at intergenic region is indicated by a “black diamond” between the two genes. B, both A72.1 and 2.6.1 – the D29-resistant mutants manifesting the highest level of resistance – sustained IS6120 integration between nucleotides G-70 and G-69 upstream of *MSMEG_1236* (*mpr*). This integration resulted in two direct repeats (indicated by black diamonds) at either sides of IS6120 at the integration site as well as introduction of the -35 promoter element “TTGACA” and a transcription factor-binding site (TFBS) for Lrp (leucine-responsive regulatory protein). Figure is drawn in a reverse orientation. Upper case letters, nucleotides introduced by IS6120 integration; underlined letters, TFBs; lower case letters, original sequence at the site; blue-font letters, direct repeats; TSS, transcription start site; atg, *mpr* start codon.

### A72.1 and 2.6.1 show elevated Mpr expression

The two mutants A72.1 and 2.6.1 sustained two transposition events each. ISMsm1 integrated in the intergenic region between *MSMEG_2105* and *MSMEG_2104* in A72.1, but integrated in *MSMEG_5393* in 2.6.1. However, in both strains, IS6120 integrated within the 168 bp intergenic region between *mpr* and *MSMEG_1237*, just upstream of *mpr*. Mpr-MSMEG_1237 interaction map on STRING database suggests possible co-expression (Figure 4A). We therefore quantified the expression of *mpr* and *MSMEG_1237* to determine if any of the genes had been affected by IS6120 integration, using Ms^Wt^ and B6.1 as control strains. Both A72.1 and 2.6.1 appear to manifest highly elevated Mpr expression, whereas *MSMEG_1237* expression remained similar among the strains (Figure 4B). We also simultaneously quantified mRNA levels at the 168 bp intergenic region (*MSGEG_1236-1237IR*) in both A72.1 and 2.6.1 as well as the control strains to detect any variations. The mRNA levels at this region don’t seem to differ much, and identical DNA sizes were seen on agarose gel for the four strains after qPCR amplification (data not shown).

**Figure 4.**
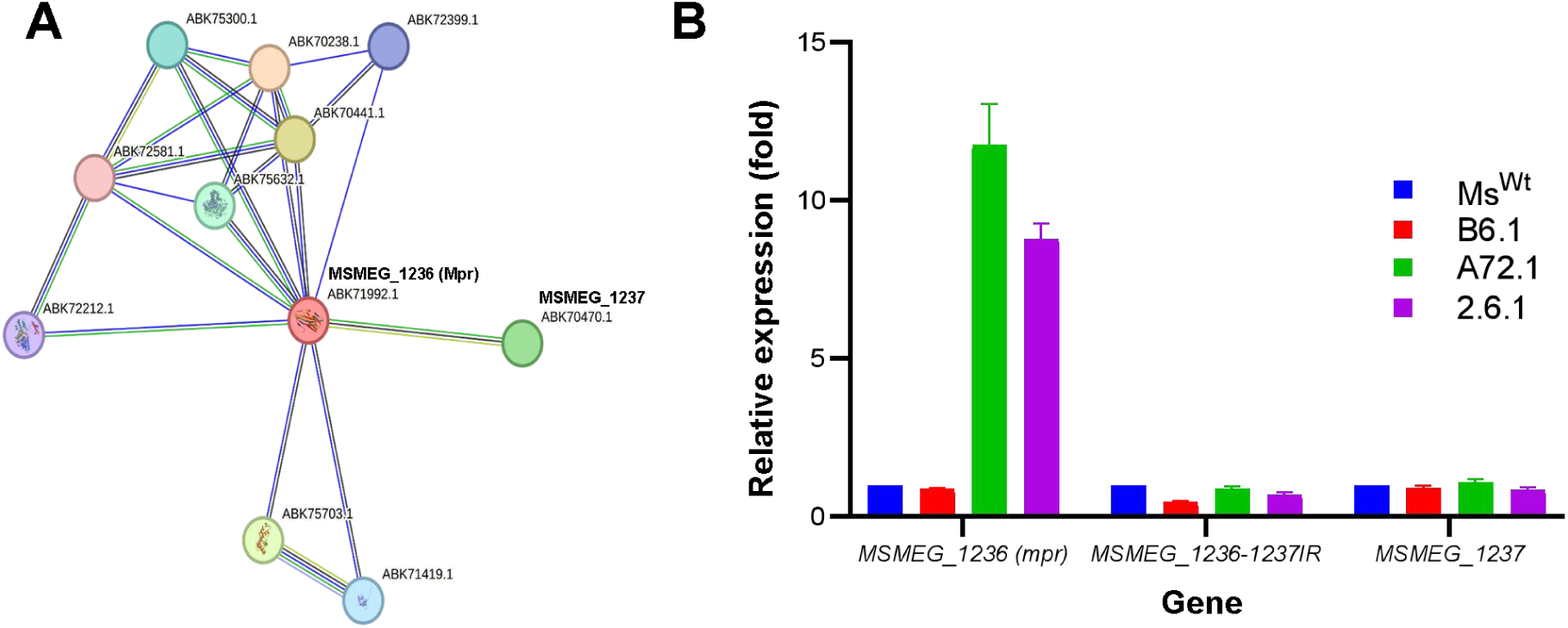
Elevated Mpr expression in A72.1 and 2.6.1. A, STRING protein-protein interaction network for Mpr. Mpr and MSMEG_1237 are adjacent to each other, and a possible co-expression is predicted in their interaction profile. B, RT-qPCR of Mpr, MSMEG_1237 as well as their intergenic region (IR) reveals that Mpr expression is elevated in A72.1 and 2.6.1.

### IS6120 integration upstream of *mpr* introduces a putative transcription factor-binding site (TFBS) and a canonical -35 promoter element

There is a 168 bp region between *mpr* and *MSMEG_1237*, and integration of IS6120 at this region appears to primarily affect the expression of *mpr*, not *MSMEG_1237* (Figure 4). In addition, IS6120 has the same orientation as *mpr* at the integration site, which raises the question of whether IS6120 integration has had some regulatory influence over *mpr*. One way this could happen is by promoter reconstitution, which inspired us to analyze the intergenic sequence prior to IS6120 integration (or <u>w</u>ild type <u>seq</u>uence, *wtseq*) and that after integration (or <u>r</u>e<u>c</u>onstituted <u>seq</u>uence, *rcseq*) for possible presence of new promoter/regulatory element(s). We used four promoter prediction programs, and each predicted at least one complete promoter in *rcseq*. One program, Sima70Pred, predicted three promoters in *wtseq* and fourteen in *rcseq*, but the remaining programs predicted no promoters in *wtseq* (Table S3). All promoters predicted by all programs in *rcseq* have the sequence “**<u>TTGACA</u>AT**TCCTCgtcccgag<u>ttgtacgtt</u>” in common, which contains a putative TFBS for leucine-responsive regulatory protein (Lrp) as well as -35 and -10 promoter elements (Figure 5). The TFBS and the canonical -35 promoter element were introduced at the site by IS6120 integration, implying that integration of IS6120 at the site brought in these two elements to reconstitute a more complete promoter than that in *wtseq*.

**Figure 5.**
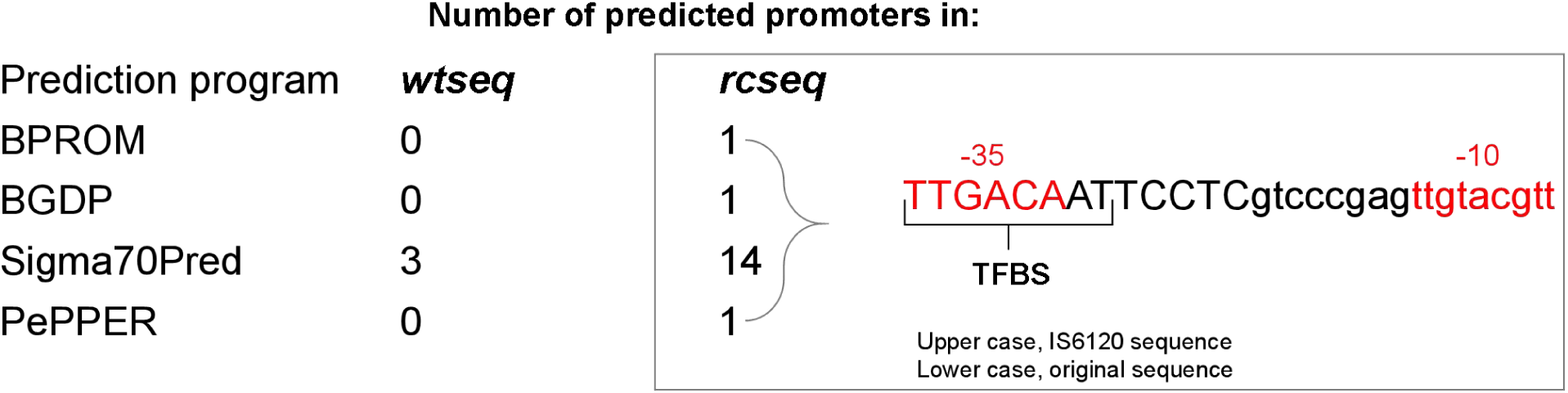
Promoter prediction in *rcseq.* Four promoter prediction programs detected the presence of a new promoter in *rcseq*. All these promoters share one sequence in common, which contains a putative TFBS, -35 and -10 promoter elements.

### IS6120 integration reconstitutes a stronger promoter upstream of *mpr*

To verify the activity of the re<u>c</u>onstituted <u>p</u>romoter (*rcp*), we carried out GFP reporter assays by constructing GFP-overexpressing strains using *rcp* or *wtp* (<u>w</u>ild type <u>p</u>romoter). We constructed Ms*^wtp^*^-GFP^ by transforming Ms^Wt^ with pMV-*wtp*-GFP, a GFP overexpression vector in which *hsp60* promoter was replaced by *wtp* (the entire 168 bp region between *mpr* and *MSMEG_1237*, amplified from Ms^Wt^). Similarly, we constructed Ms*^rcp^*^-GFP^ in which the *hsp60* promoter was replaced by *rcp* (a region of equal length directly upstream of *mpr*, amplified from A72.1). Ms^GFP^, a strain overexpressing GFP from the *hsp60* promoter, was used as a control.

The first evidence of promoter activity manifested at colony level after transformation of bacteria with the respective vectors. Ms^GFP^ and Ms*^rcp^*^-GFP^ colonies manifested the characteristic green color of GFP. Lawn of Ms*^rcp^*^-GFP^ also shows the characteristic green coloration on plain 7H10 plates (Figure 6A). Though Ms*^rcp^*^-GFP^ produces lower live fluorescence than Ms^GFP^, it produces far greater fluorescence than Ms*^wtp^*^-GFP^, suggesting that introduction of the new elements by IS6120 significantly increased the strength of the promoter at the integration site (Figure 6B). We also counted GFP events for each of the strains by flow cytometry, and the number of GFP events detected for Ms*^rcp^*^-GFP^ was clearly far greater than that for Ms*^wtp^*^-GFP^, which constitutes further evidence of superior promoter activity (Figure 7). Finally, confocal micrographs corroborate the flow cytometry data, showing that *wtp* drives significantly lower GFP expression than *rcp*, and that the GFP signals produced by Ms*^rcp^*^-GFP^ and Ms^GFP^ are somewhat identical (Figure 8). Collectively, these imply that IS6120 integration upstream of *mpr* brings about replacement of *wtp* by *rcp*, which raises the question of whether *M. smegmatis* regulates the expression of Mpr by promoter reconstitution. In addition, we attempted to overexpress Mpr in Ms^Wt^ by constructing Ms*^wtp^*^-Mpr^ and Ms*^rcp^*^-Mpr^ using the vectors pMV-*wtp*-*mpr* or pMV-*rcp*-*mpr* respectively. However, strains transformed with pMV-*rcp*-*mpr* rarely formed colonies on the plate (Figure S2). Due to low efficiency of Ms*^rcp^*^-Mpr^ to form colonies, we are unable to provide any definitive inference *vis-à-vis* the relationship between *rcp* and Mpr expression or D29 resistance. However, it does appear like expressing Mpr using *rcp* exerts toxicity on the bacterial cells.

**Figure 6.**
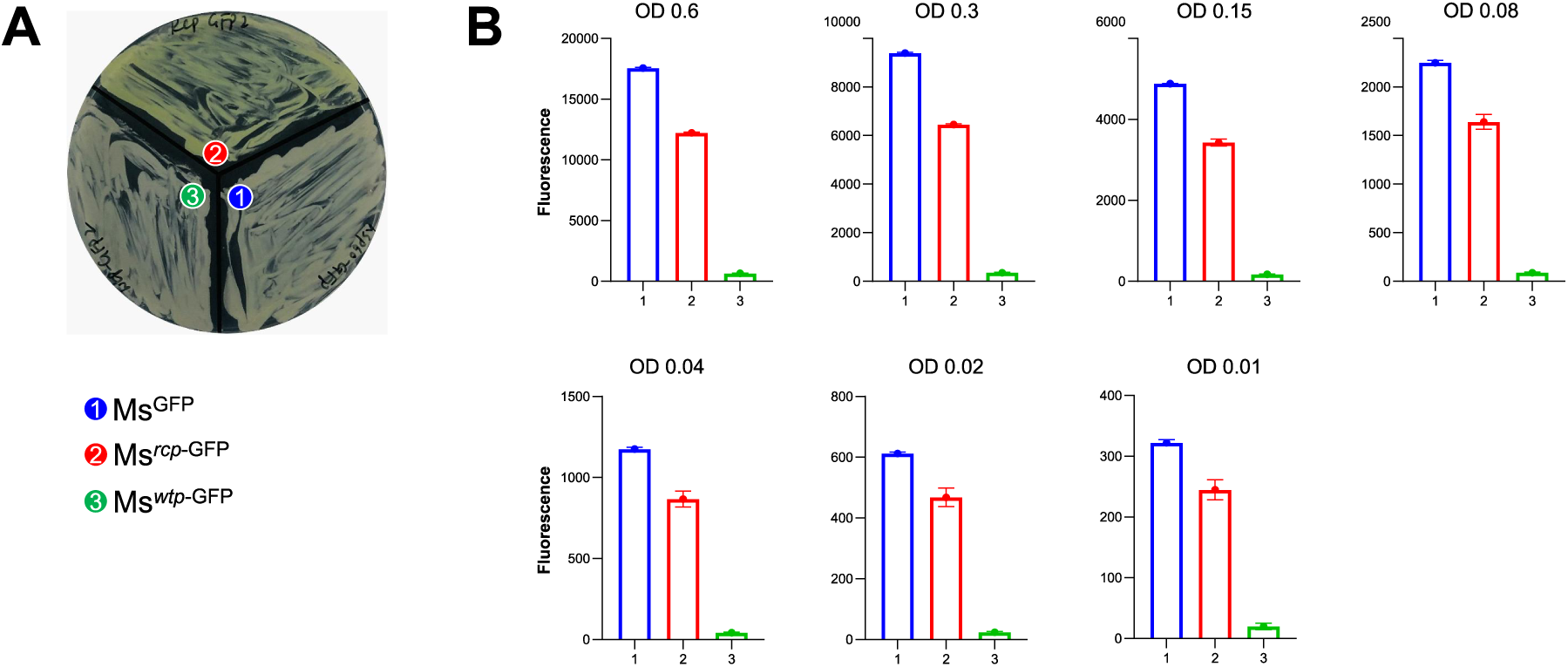
Quantification of *wtp*- and *rcp*-driven green fluorescence. A, Lawn appearance of GFP-expressing strains using *hsp60*, *wtp* and *rcp*. Ms*^rcp^*^-GFP^ shows the characteristic green coloration of GFP. B, Live fluorescence of each strain as quantified using a microplate reader at different optical densities indicates that *rcp* drives higher fluorescence than *wtp*.

**Figure 7.**
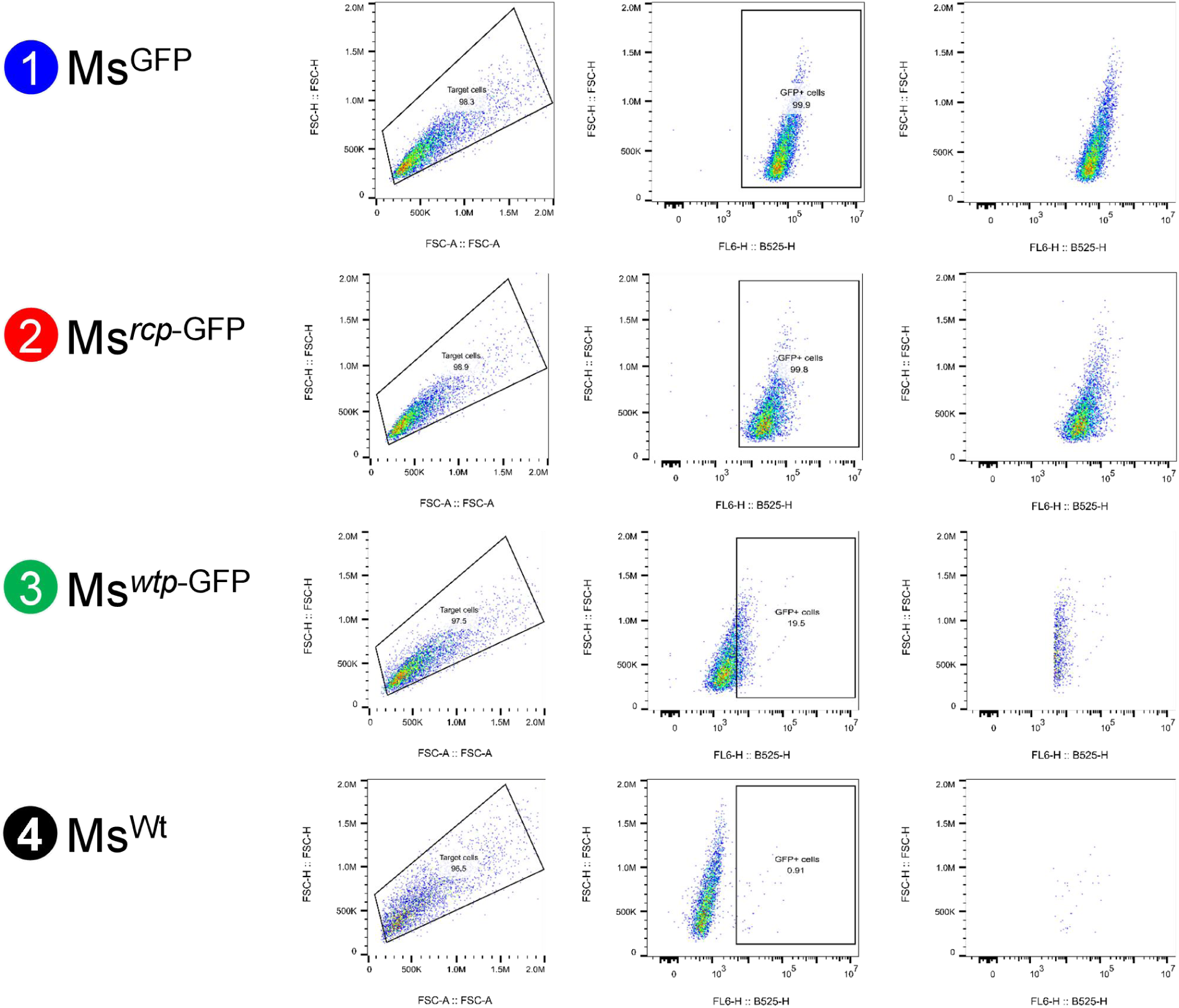
Quantification of *wtp*- and *rcp*-driven GFP events by flow cytometry. Quantification of GFP events places *hsp60* and *rcp* on par with each other, whereas *wtp* falls far behind in comparison. Quantification was carried out in a 10,000-cell population for each strain. Populations expressing GFP using *hsp60* (Ms^GFP^), *rcp* (Ms*^rcp^*^-GFP^) and *wtp* (Ms*^wtp^*^-GFP^) had 99.9, 99.8 and 19.5 % GFP-positive cells respectively. In contrast, the negative control (Ms^Wt^ population) had a negligible 0.91 % GFP-false positive cells.

**Figure 8.**
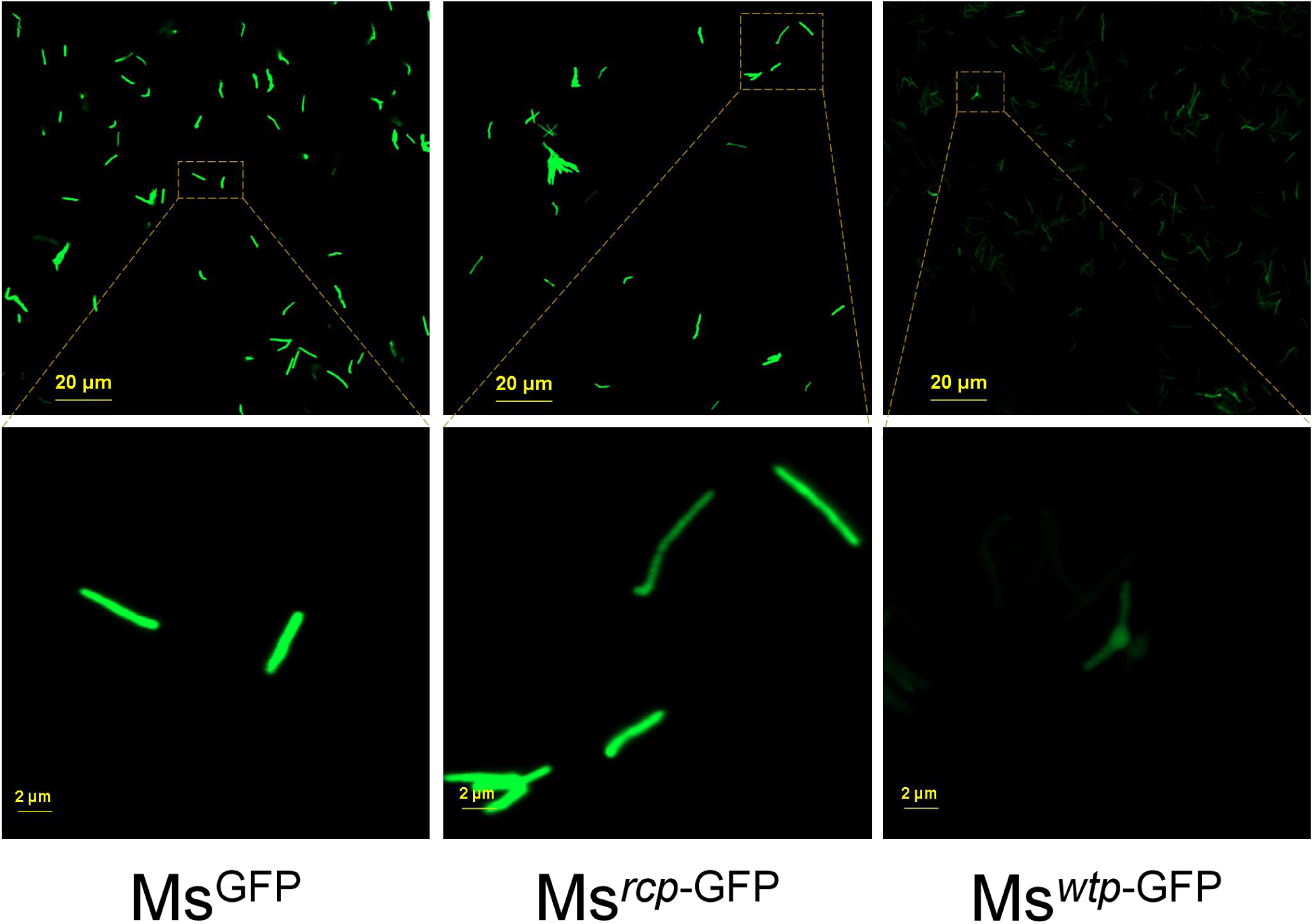
Quantification of fluorescence signal. Fluorescence signal driven by *hsp60*, *rcp* and *wtp* were quantified by confocal laser scanning microscopy. Strength of signal produced by *hsp60* and *rcp* appear to be similar, whereas *wtp* drives far weaker fluorescence signal under the microscope.

### D29 resistance had no effect on *M. smegmatis* resistance to selected drugs

To assess potential fitness tradeoffs associated with D29 resistance, we performed drug susceptibility testing (DST) on Ms^Wt^ and three resistant mutants (B6.1, A72.1, 2.6.1) against seven frontline anti-tubercular agents: isoniazid, rifampicin, linezolid, clofazimine, bedaquiline, levofloxacin, and ethambutol. Remarkably, all strains—including the resistant mutants—exhibited identical susceptibility profiles to these antibiotics (Table S4). This lack of divergence suggests that D29 resistance does not impinge upon core cellular processes targeted by these drugs, nor does it incur detectable collateral costs in tolerance to these drugs. These findings imply that mycobacterial adaptation to phage pressure may occur independently of classical antibiotic resistance pathways.

## Discussion

AMR triggered a renewed interest in phage therapy as a potential alternative to antibiotic therapy (3,33,34). However, phage resistance represents a potent threat to sustainable usability of phage therapy, addressing which would require dissecting the mechanisms of phage interactions with their bacterial hosts. Efforts towards dissecting phage-bacteria interactions revealed invaluable details regarding receptor requirements (19) and adaptive responses to phage pressure, including the roles played by IS elements (20,42,43) as well as Mpr in M. smegmatis (15). Studies on Mpr role in D29 resistance have relied on vector-borne overexpression of the gene. Here, we report two D29-resistant mutants with elevated Mpr expression that likely stemmed from promoter reconstitution by IS6120 integration directly upstream of *mpr*.

ISs are mobile genetic elements that are widespread in bacterial genomes and play important roles in bacterial evolution and adaptation to stress. It is becoming increasingly evident that phage infection triggers IS transposition, which causes these elements to integrate into certain genes in bacteria and cause phage resistance via different mechanisms (20,42,43). For example, TM4 infection of *M. smegmatis* triggers IS1096 transposition and integration into *lsr2*, hence activating the LOS cluster, causing PIMs accumulation and reduced phage adsorption (20). Such transposition events often cause large insertions that, coupled with widespread endogenous presence of IS elements in the chromosome, may often be missed by mere SNP calling. This explains why they were not detected during the initial whole-genome sequence analysis. Therefore, we analyzed the raw sequence data for all sequenced D29-resistant mutants using ISMapper (27), and we detected several IS transposition events (Figure 3A). We investigated one particular transposition event in A72.1 and 2.6.1, which resulted in integration of IS6120 within the 168 bp region between *MSMEG_1236* (<u>m</u>ulti-copy <u>p</u>hage resistance gene, *mpr*) and *MSMEG_1237* (Figure 3B). Mpr is a membrane-bound DNA exonuclease that, upon overexpression, defends *M. smegmatis* against D29 by cleaving phage DNA following its injection into the cytoplasm (14–16). Mpr is required for the emergence of spontaneous D29-resistant mutants, and such role does not require any mutation in the gene but overexpression of the wild type gene. However, the genetic basis for emergence of D29-resistant mutants with elevated Mpr expression remained elusive. Here, we uncovered that both A72.1 and 2.6.1 have IS6120 integration directly upstream of *mpr*, and both show profoundly elevated expression of Mpr, not MSMEG_1237 (Figure 4B). In addition, the integrated IS6120 element runs in the same orientation (5’-3’) as *mpr*, not *MSMEG_1237*, which could partly explain the IS transposition effect on Mpr rather than MSMEG_1237. Further analysis of the sequence at the integration site revealed that IS6120 introduced a -35 promoter element and a TFBS for Lrp, which were not originally present at the site, hence reconstituting a new, fuller promoter (Figure 5). Promoter reporter assays reveal that the reconstituted promoter (*rcp*) has activity that is far superior than that of the original promoter (*wtp*) and is nearly as good as the BCG promoter *hsp60* (Figures 7 and 8). This certainly begs the question of whether elevated Mpr expression in spontaneous D29-resistant mutants has always been due to reconstitution of a fuller, more powerful promoter upstream of *mpr*, which may also explain why emergence of such mutants require the presence of an intact *mpr* (15). Failure to isolate recombinant strains overexpressing *rcp-mpr* suggest extreme toxicity, and this is consistent with previous effort to express *mpr* in *M. smegmatis* using the *hsp60* promoter (14). This mechanism of gene regulation by promoter reconstitution is common in bacteria (44–46), and the extreme toxicity may well be reminiscent of such situations where inherent protein toxicity often makes it difficult to study (10,45). Mechanism of Mpr-mediated defense is believed to be non-abortive in nature (16), and it remains to be seen whether A72.1 and 2.6.1 were able to grow normally in the face of elevated Mpr expression because the observed expression levels were tolerable, or possibly due to neutralization of toxicity by other factors, likely the second transposition event or other mutations detected in each of these strains. DEMs of D29 have been reported for vector-borne Mpr-overexpressing *M. smegmatis* (16), but we were unable to isolate a DEM for A72.1 and 2.6.1 in this study. It is not clear whether this could be due to chromosome-borne, *rcp*-driven Mpr overexpression or absolute resistance in these strains (13).

Mpr contains two major domains - a transmembrane domain (amino acids 33-50) and the domain of unknown function DUF4352 (amino acids 83-211). DUF4352 is believed to be of viral origin, suggesting that the bacteria likely acquired it during the course of evolution with viral predators, then kept it for anti-viral defense (16). The fact that Ms^Wt^ is susceptible to D29 is evidence that wild type level of Mpr expression does not confer resistance. However, regulation of Mpr expression by promoter reconstitution could suggest that the gene is most useful to the bacteria under D29-induced stress, not normal conditions. From our observations, Mpr appears to be the primary anti-D29 resistance tool in *M. smegmatis*.

Events such as gene amplification-duplication are transient adaptive responses to increase bacterial fitness, and are usually reversible due to the high fitness costs they exert (47,48). IS transposition could also be reversible (43), but we haven’t seen any evidence of such reversion in A72.1 and 2.6.1 despite high expression of the toxic protein Mpr, and it remains to be seen whether this might happen in later generations of the bacteria. Tolerance of A72.1 and 2.6.1 to drugs appears to be unaffected, meaning that the strains had not traded off tolerance to the drugs (at least those tested in this study) for D29 resistance. They also fare very well under normal growth conditions, indicating lack of growth-related fitness cost associated with D29 resistance in the strains.

This study reveals a form of gene regulation by promoter reconstitution in *M. smegmatis*. We report that exposure of *M. smegmatis* to the lytic mycobacteriophage D29 triggers the transposition and integration of IS6120 directly upstream of *mpr*, resulting in reconstitution of a fuller, stronger promoter. This likely represents a previously elusive factor behind emergence of D29-resistant mutants with elevated Mpr expression.

## Acknowledgments

This study was supported by the National Key R&D Program of China (2021YFA1300900), the Major Project of Guangzhou National Laboratory (GZNL2024A01009, GZNL2025C01003), the National Natural Science Foundation of China (82502762).

## Data availability

The next generation sequencing reads for Ms^Wt^, 2.6.1 and A72.1 have been deposited at the NCBI Sequence Read Archive, and can be accessed with SAMN55349604, SAMN55349605 and SAMN55349614 respectively.

