## Supplemental files for "Likely role of promoter reconstitution in Mpr-mediated D29 resistance by *Mycobacterium smegmatis*"

**Table S1** Different IS elements, their segments and sizes in *M. smegmatis*

**Figure S1** Prevalence of IS elements in *M. smegmatis* genome.

**Table S2** IS rearrangements in the D29-resistant mutants

**Table S3** Promoter prediction in wild type and reconstituted sequences upstream of *mpr*

**Figure S2** Colony formation on media plates by Mpr-overexpressing strains.

**Table S1 Different IS elements, their segments and sizes in *M. smegmatis***

| IS element | Segments | Size (ISFinder) |
|---|---|---|
| IS1096 | *tnpA* and *tnpR* | 2259 bp |
| IS1137 | *orfA* and *orfB* | 1361 bp |
| IS1549 | - | 1640 bp |
| IS6120 | - | 1486 bp |
| ISMsm1 | *orfA* and *orfB* | 1345 bp |
| ISMsm2 | - | 1126 bp |
| ISMsm3 | - | 1695 bp |
| ISMsm4 | - | 1458 bp |
| ISMsm5 | - | 1103 bp |
| ISMsm6 | - | 1765 bp |
| ISMsm7 | *orfA* and *orfB* | 1280 bp |
| ISMsm8 | - | 1457 bp |
| ISMsm9 | - | 1093 bp |
| ISMsm10 | - | 1661 bp |
| ISMsm11 | - | 1658 bp |
| ISMsm12 | - | 3256 bp |

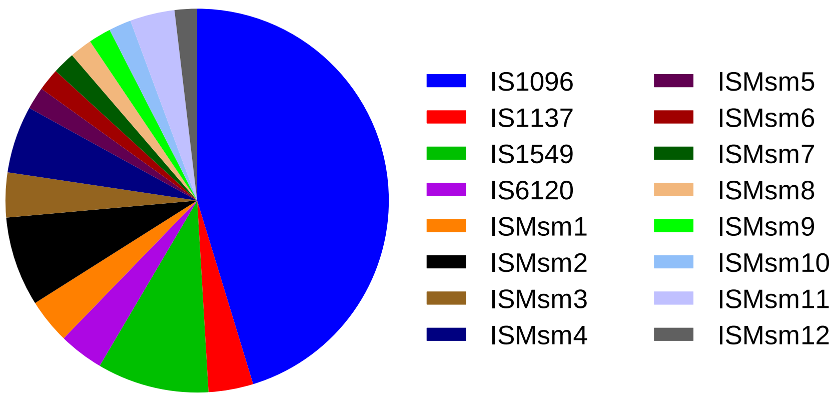


**Figure S1 Prevalence of IS elements in *M. smegmatis* genome.** The *M. smegmatis* genome encodes 16 known IS elements. The most prevalent IS element encoded by the genome is IS1096, followed by IS1549, ISMsm2 and ISMsm4.

**Table S2 IS rearrangements in the D29-resistant mutants**

| Insertion position | IS element | Region | Affected gene(s) | Product |
|---|---|---|---|---|
| 3822918-3822926 | IS1096 | Coding | *MSMEG_3756 (rrlA)* | 23S ribosomal RNA |
| 1307414-1307406 | IS6120 | Intergenic | *MSMEG_1236* ♦ *MSMEG_1237* | Mpr protein and a hypothetical protein, respectively |
| 1822025-1822016 | IS6120 | Coding | *MSMEG_1730* | ISMsm1, transposase *orfB* |
| 3403974-3403965 | IS6120 | Coding | *MSMEG_3324* | Hypothetical protein |
| 4817559-4817569 | IS6120 | Coding | *MSMEG_4724* | Oligoribonuclease |
| 5192312-5192302 | IS6120 | Coding | *MSMEG_5091* | Hypothetical protein |
| 2181906-2181903 | ISMsm1 | Intergenic | *MSMEG_2105* ♦ *MSMEG_2104* | ATP-dependent DNA ligase and a GntR-family protein transcriptional regulator, respectively |
| 3358211-3358215 | ISMsm1 | Intergenic | *MSMEG_3275* ♦ *MSMEG_3276* | RNA polymerase sigma factor (sigma-70 family protein) and an integral membrane protein, respectively |
| 5471133-5471130 | ISMsm1 | Coding | *MSMEG_5393 (kdpB)* | K+ - transporting ATPase, B subunit |

**Table S3 Promoter prediction in wild type and reconstituted sequences upstream of *mpr***

| Prediction program | Promoter sequence(s) in: | |
|---|---|---|
| | **Wild type sequence (*wtseq*)** | **Reconstituted sequence (*rcseq*)** |
| BPROM | - | -35 element = TTGACA<br>-10 element = ttgtacgtt<br>TFBS (Lrp) = TTGACAAT |
| BDGP | - | AGGTATTGACAATTCCTCgtcccgagttgtacgttcgcctgttcgcctgg [*] |
| Sigma70Pred | 1. CGGCACGACCCGACCAAGGACAGGCACGAGGTTGTCGGTCCCGAGTTGTACGTTCGCCTGTTCGCCTGGCATGGGGCCCGG<br>2. CACTACCTCCTGATTTGAGTGAGCCGCACAGCTTCGCATGGCCCACCGACAAATCGTCGCGACGGCACGACCCGACCAAGG [#]<br>3. ACAGGCACGAGGTTGTCGGTCCCGAGTTGTACGTTCGCCTGTTCGCCTGGCATGGGGCCCGGCGCAAGACATGAAGGAACG | 1. AGGTATTGACAATTCCTCGTCCCGAGTTGTACGTTCGCCTGTTCGCCTGGCATGGGGCCCGGCGCAAGACATGAAGGAACG<br>2. AGCACGCCGGAACGGAGGTCGCCTGAAACACCCCGATCCACAGGTATTGACAATTCCTCGTCCCGAGTTGTACGTTCGCCT<br>3. ATTGACAATTCCTCGtcccgagttgtacgttcgcctgttcgcctggcatggggcccggcgcaagacatgaaggaacgatca<br>4. CCTGAAACACCCCGATCCACAGGTATTGACAATTCCTCGTCCCGAGTTGTACGTTCGCCTGTTCGCCTGGCATGGGGCCCG<br>5. CGCCTGAAACACCCCGATCCACAGGTATTGACAATTCCTCGTCCCGAGTTGTACGTTCGCCTGTTCGCCTGGCATGGGGCC<br>6. CGGTCAGCACGCCGGAACGGAGGTCGCCTGAAACACCCCGATCCACAGGTATTGACAATTCCTCGTCCCGAGTTGTACGTT<br>7. CTGAAACACCCCGATCCACAGGTATTGACAATTCCTCGTCCCGAGTTGTACGTTCGCCTGTTCGCCTGGCATGGGGCCCGG<br>8. GCACGCCGGAACGGAGGTCGCCTGAAACACCCCGATCCACAGGTATTGACAATTCCTCGTCCCGAGTTGTACGTTCGCCTG<br>9. GCCTGAAACACCCCGATCCACAGGTATTGACAATTCCTCGTCCCGAGTTGTACGTTCGCCTGTTCGCCTGGCATGGGGCCC |

| | | |
|---|---|---|
| | | 10. GGTATTGACAATTCCTCGTCCCGAGTTGTACGTTCGCCTGTTCGCCTGGCATGGGGCCCGGCGCAAGACATGAAGGAACGA<br>11. GGTCAGCACGCCGGAACGGAGGTCGCCTGAAACACCCCGATCCACAGGTATTGACAATTCCTCGTCCCGAGTTGTACGTTC<br>12. GTATTGACAATTCCTCGTCCCGAGTTGTACGTTCGCCTGTTCGCCTGGCATGGGGCCCGGCGCAAGACATGAAGGAACGAT<br>13. GTCGCCTGAAACACCCCGATCCACAGGTATTGACAATTCCTCGTCCCGAGTTGTACGTTCGCCTGTTCGCCTGGCATGGGG<br>14. TATTGACAATTCCTCGTCCCGAGTTGTACGTTCGCCTGTTCGCCTGGCATGGGGCCCGGCGCAAGACATGAAGGAACGATC |
| PePPER | - | TTGACAATTCCTCgtcccgagttgtacgtt |

BPROM, bacterial promoter prediction program; TFBS, transcription factor-binding site; Lrp, leucine-responsive regulatory protein; BDGP, Berkeley drosophila genome project; PePPER, prediction of prokaryote promoter elements and regulons.

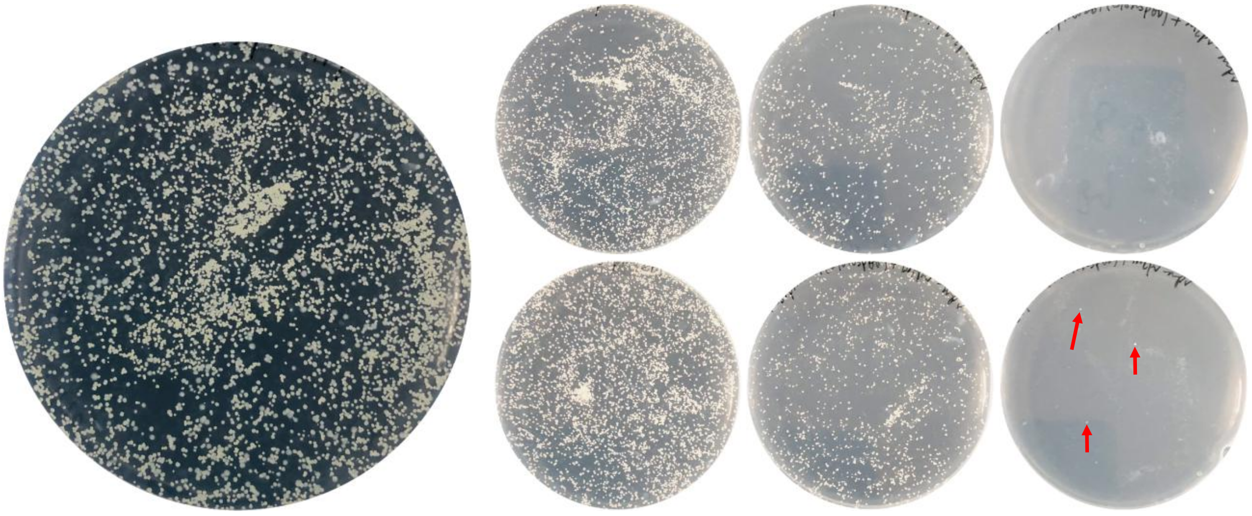


**Figure S2 Colony formation on media plates by Mpr-overexpressing strains.** Control strains (Ms$^{GFP}$ and Ms$^{Wt}$:pMV261) and Ms$^{wtp\text{-}mpr}$ efficiently formed colonies on 7H10 plates after transformation. Ms$^{rcp\text{-}mpr}$ on the other hand rarely formed colonies on the plates. The few colonies formed by Ms$^{rcp\text{-}mpr}$ are indicated by red arrows.

**Table S4 Drug susceptibility profiles of selected D29-resistant mutants**

| | Drugs/MIC (μg/mL) | | | | | | |
|---|---|---|---|---|---|---|---|
| **Strains** | **INH** | **RIF** | **LZD** | **CLF** | **BDQ** | **LEV** | **EMB** |
| Ms$^{Wt}$ | 16 | >16 | 2 | 1 | 0.03125 | 0.25 | 1 |
| B6.1 | 16 | >16 | 2 | 1 | 0.03125 | 0.25 | 1 |
| A72.1 | 16 | >16 | 2 | 1 | 0.03125 | 0.25 | 1 |
| 2.6.1 | 16 | >16 | 2 | 1 | 0.03125 | 0.25 | 1 |
